# Decoding of selective attention to continuous speech from the human auditory brainstem response

**DOI:** 10.1101/259853

**Authors:** Octave Etard, Mikolaj Kegler, Chananel Braiman, Antonio Elia Forte, Tobias Reichenbach

## Abstract

Humans are highly skilled at analysing complex acoustic scenes. The segregation of different acoustic streams and the formation of corresponding neural representations is mostly attributed to the auditory cortex. Decoding of selective attention from neuroimaging has therefore focussed on cortical responses to sound. However, the auditory brainstem response to speech is modulated by selective attention as well, as recently shown through measuring the brainstem’s response to running speech. Although the response of the auditory brainstem has a smaller magnitude than that of the auditory cortex, it occurs at much higher frequencies and therefore has a higher information rate. Here we develop statistical models for extracting the brainstem response from multi-channel scalp recordings and for analysing the attentional modulation according to the focus of attention. We demonstrate that the attentional modulation of the brainstem response to speech can be employed to decode the attentional focus of a listener from short measurements of ten seconds or less in duration. The decoding remains accurate when obtained from three EEG channels only. We further show how out-of-the-box decoding that employs subject-independent models, as well as decoding that is independent of the specific attended speaker is capable of achieving similar accuracy. These results open up new avenues for investigating the neural mechanisms for selective attention in the brainstem and for developing efficient auditory brain-computer interfaces.

## Introduction

Humans have an extraordinary capability to analyse crowded auditory scenes. We can, for instance, focus our attention on one of two competing speakers and understand her or him despite the distractor voice (Middlebrooks et al., 2017). People with hearing impairment such as sensorineural hearing loss, however, face major difficulty with understanding speech in noisy environments, and this difficulty persists even when they wear auditory prosthesis such as hearing aids or cochlear implants (Armstrong et al., 1997). Auditory prosthesis could potentially aid with understanding speech in noise through selectively enhancing a target speech, for instance based on its direction, using algorithms such as beam forming (Kidd et al., 2015). However, such selective enhancement requires knowledge of which sound the user aims to attend to. Current research therefore attempts to decode an individual’s focus of selective attention to sound from non-invasive brain recordings (O’Sullivan et al., 2014; Mirkovic et al., 2015; Biesmans et al., 2016; Fuglsang et al., 2017). If such decoding worked in real time, it could inform the sound processing in an auditory prosthesis. It could also form the basis of a non-invasive brain-computer interface for motor-impaired patients with brain injury, for instance, who may not be able to respond behaviourally. Moreover, such decoding of selective attention could be employed clinically for a better understanding and characterization of hearing loss.

Neural activity in the cerebral cortex, especially in the delta (1 – 4 Hz) and theta (4 – 8 Hz) frequency bands, tracks the amplitude envelope of a complex auditory stimulus such as speech (Ding and Simon, 2012; Giraud and Poeppel, 2012; Power et al., 2012; Ding and Simon, 2014). The tracking is shaped by selective attention to one of several sound sources and can be measured from electrocorticography (ECoG) (Mesgarani and Chang, 2012), and noninvasively from magnetoencephalograpy (MEG) (Ding and Simon, 2012), as well as from the clinically more applicable electroencephalography (EEG) (Kerlin et al., 2010; Horton et al., 2013). Attention to one of two competing voices has been successfully decoded from single trials of one minute in duration using MEG (Ding and Simon, 2012) as well as EEG (O’Sullivan et al., 2014; Mirkovic et al., 2015; Fiedler et al., 2017). Further optimization of the involved statistical modelling led to an accurate decoding of the focus of selective attention from still shorter recordings lasting less than 30 s (Biesmans et al., 2016; Van Eyndhoven et al., 2017). Moreover, a subject’s changing focus of attention could be detected within tens of seconds from EEG data, and even faster from MEG data, when combined with additional sparse statistical modeling (Miran et al., 2018).

Recently we showed that subcortical neural activity is consistently modulated by selective attention as well (Forte et al., 2017). To this end we developed a method to measure the response of the auditory brainstem to natural non-repetitive speech. We employed empirical mode decomposition (EMD) to extract a waveform from the speech signal that, at each time instance, oscillates at the fundamental frequency of the voice. We then correlated this fundamental waveform to the neural recording obtained from a few scalp electrodes. We observed a peak in the cross-correlation at a latency of 9 ms, evidencing a neural response at the fundamental frequency with a subcortical origin. This method determined the brainstem response to the voiced parts of speech, and in particular to its pitch. When volunteers listened to two competing speakers, we observed that the brainstem response to the fundamental frequency of each speaker was larger when the speaker was attended than when she or he was ignored.

Because the brainstem response to speech that we measured occurs at the fundamental frequency of speech, typically between 100 – 300 Hz, it is ten- to hundredfold faster than the cortical tracking of the speech envelope. The rapidness of the brainstem response could imply a high information rate, despite the small magnitude of the response that is below that of cortical responses. We therefore wondered if the brainstem response to natural speech can be detected from high density EEG, that is typically used to capture the cortical activity, and whether it can be used to efficiently decode auditory attention.

## Materials and Methods

### Participants

18 healthy adult English native speakers (aged 22.8 ± 1.9 year, four females), with no history of auditory or neurological impairments participated in the study. All participants provided written informed consent. The experimental procedures were approved by the Imperial College Research Ethics Committee.

### Experimental Design and Statistical Analysis

We employed the same experimental design that we used previously to measure the brainstem response to non–repetitive speech and its modulation through selective attention (Forte et al., 2017). In particular, approximately ten-minute long continuous speech samples from a male and female speaker were obtained from publicly available audiobooks (librivox.org). For the female voice excerpts from *“The Children of Odin”* (chapters 2 and 4) and *“The Adventures of Odysseus and the Tale of Troy”* (part 2, chapter 8), all by Pádraic Colum and read by Elizabeth Klett, were selected. For the male voice excerpts from *“Tales of Troy: Ulysses the Sacker of Cities”* by Andrew Lang (section 11) and *“The Green Forest Fairy Book”* by Loretta Ellen Brady (chapter 10), all read by James K. White, were used. The first story from the female speaker was employed when presenting speech in quiet. The two other female speech samples were used to generate two stimuli with two competing speakers by mixing each with one sample from the male speaker, at equal root-mean-square amplitude.

Participants first listened to the stimulus with a single speaker without background noise. They then listened to the two stimuli with two competing speakers each. They were instructed to exclusively attend either the male or female voice in the first stimulus, and to attend to the speaker they previously ignored in the second one. Whether a subject was instructed to first attend the male speaker and then the female speaker or *vice versa* was determined randomly for each subject. Each stimulus was presented in four parts of approximately equal duration (~2.5 minutes), and comprehension questions were asked after each part. All stimuli were delivered diotically, that is, the same waveforms were delivered to the right and left ears, at 76 dB(A) SPL (A-weighted frequency response) using Etymotic ER-3C insert tube earphones to minimise artifacts. The sound intensity was calibrated with an ear simulator (Type 4157, Brüel & Kjaer, Denmark). EEG recordings were obtained during the stimuli and their statistical analysis was performed using custom Matlab and Python code and functions from the MNE toolbox (Gramfort et al., 2013; Gramfort et al., 2014) as described below.

### Neural data acquisition and processing

Neural activity was recorded at 1 kHz through a 64-channel scalp EEG system using active electrodes (actiCAP, BrainProducts, Germany) and a multi-channel EEG amplifier (actiCHamp, BrainProducts, Germany). The electrodes were positioned according to the standard 10-20 system and referenced to the right earlobe. The EEG recordings were band-pass filtered offline between 100 and 300 Hz (low pass: linear phase FIR filter, cutoff (−6 dB) 325 Hz, transition bandwidth 50 Hz, order 66; high pass: linear phase FIR filter, cutoff (−6 dB) 95 Hz, transition bandwidth 10 Hz, order 364; both: one-pass forward and compensated for delay) and then referenced to the average. When only using three channels for the decoding, all channels except the two mastoids TP9 and TP10 and the vertex Cz were discarded, and the filters described above were applied. The audio signals were simultaneously recorded by the amplifier at a sampling rate of 1 kHz through an acoustic adapter (Acoustical Stimulator Adapter and StimTrak, BrainProducts, Germany), and were used to align the neural responses to the stimuli. A 1 ms delay of the acoustic signal introduced by the earphones was taken into account by shifting the audio signal forward by 1 ms with respect to the neural response.

### Computation of the fundamental waveform of speech

We employed Empirical Mode Decomposition (EMD) to extract a waveform from each speech signal that, at each time instance, oscillates at the fundamental frequency of the voice; we refer to it as the fundamental waveform (Forte et al., 2017). EMD is indeed well suited to analyze data that results from non-stationary and nonlinear processes such as speech production, and has been successfully used for pitch detection (Huang and Pan, 2006). The fundamental waveform was downsampled to 1 kHz, the sampling rate of the neural recordings, and filtered between 100 and 300 Hz as described above. Silent or unvoiced parts of the speech produced some segments where the fundamental waveform was equal to zero. For the stimuli with a single speaker, we excluded such segments from the further analysis. For the stimuli with two competing speakers we excluded the few segments where the fundamental waveform of one of the two voices was entirely zero as attention could not be decoded in this case.

We also computed a proxy of the fundamental waveform by band-pass filtering the audio signal in the range of the fundamental frequency. We thereby employed FIR filters with corner frequencies of 100 Hz and 200 Hz for the male voice (linear-phase FIR filter, lower cutoff (−6 dB): 90 Hz, transition bandwidth 17.5 Hz, higher cutoff (−6 dB): 210 Hz, transition bandwidth 17.5 Hz, order 237, one pass forward and compensated for delay), as well as corner frequencies of 150 Hz and 250 Hz for the female voice (linear-phase FIR filter, lower cutoff (−6 dB): 135 Hz, transition bandwidth 25 Hz; higher cutoff (−6 dB): 275 Hz, transition bandwidth 25 Hz, order 157, one pass forward and compensated for delay). We employed the band-pass filtered audio signals to obtain the results on attention reported in Figure 7-B. All other results presented here were obtained from waveforms extracted by EMD.

### Backward model

We first used a linear spatio-temporal backward model to reconstruct the fundamental waveform of speech from the neural recordings. Specifically, at each time instance *t_n_*, the fundamental waveform *y*(*t_n_*) was estimated as a linear combination of the neural recordings *x_j_* (*t_n_* + *τ_k_*) as well as their Hilbert transform 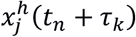 at a delay *τ_k_*:

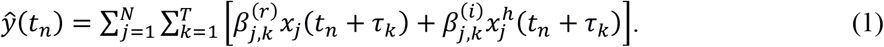

The index *j* refers hereby to the recording channel, and 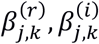 are a set of real coefficients to determine. We used a set of *T* = 25 possible delays *τ_k_* ranging from −5 ms to 19 ms with an increment of 1 ms. The Hilbert transform of each recording channel was included in Equation (1), denoted with the upper index *h*, to allow the reconstruction of the fundamental waveform from these signals as well. The Hilbert transform of a sinusoid results in a phase shift of π/2, which equates to a temporal shift of a quarter period. Even narrow-band signals such as our band-pass filtered EEG recordings contain, however, a range of frequencies. While the Hilbert transform of these signals can still be interpreted as a phase shift of π/2, it can no longer be obtained by a temporal shift. The Hilbert transforms therefore add another set of predictors in Equation (1) that are independent of the time-shifted EEG signals, and that thereby aid the reconstruction of the fundamental waveform.

The model’s coefficients can be assembled into complex coefficients 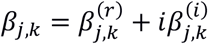 that encode accordingly the amplitude of the brainstem response, the temporal delay as well as the phase difference between stimulus and response. We thus obtained *T* = 25 temporal delays that, together with the *N*=64 recording channels, led to 1,600 complex model coefficients.

The model coefficients were then estimated for each subject using a regularised ridge regression as *β* = (*X^t^X* + *λI*)^−1^*X^t^y*, in which *X* is the design matrix of dimension *n_p_* × 2*NT* with *n_p_* the number of samples available in the recording, and *λ* is a regularisation parameter (Hastie et al., 2009). In particular, the columns of the design matrix are the neural recordings *x_j_*(*t_n_* + *τ_k_*) at the different time points *t_n_* as well as their Hilbert transforms 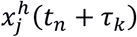. To normalise for differences between datasets, *λ* can be written as *λ* = *λ_n_ e_m_* where *e_m_* is the mean eigenvalue of *X^t^X* and *λ_n_* is a normalised regularisation coefficient (Biesmans et al., 2016).

A five-fold cross-validation procedure was implemented to evaluate the model. In each of five iterations, and for each participant, four folds of the ten-minute data were used to compute the model coefficients, yielding about eight minutes of training data. The remaining fifth fold, two minutes of testing data, served to estimate the fundamental waveform and to compute the performance of the model. The performance was quantified by dividing the reconstructed 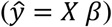 and the actual (*y*) fundamental waveforms obtained on the testing data in ten-seconds long segments and computing Pearson’s correlation coefficient between these waveforms for each segment. The correlation values obtained over the five testing folds were pooled to determine the mean and standard error of the reconstruction performance. This performance was determined for 50 different normalised regularization parameters with values ranging from 10^−15^ to 10^15^. For each subject, the regularization parameter that yielded the largest reconstruction performance was chosen as the optimal regularization parameter.

The procedure above, including the use of the Hilbert transform of the EEG data, was employed both when reconstructing the fundamental waveform obtained from EMD as well as when estimating the fundamental waveform obtained from band-pass filtering the speech signal (see below).

The Python code for computing the complex coefficients of the backward model, together with a sample of a fundamental waveform and the corresponding EEG recordings, is on Github (Kegler et al.).

### Significance of the fundamental waveform reconstruction

To determine if the linear backward models showed a significant brainstem response to the fundamental frequency, we also computed, for each subject, one noise model as a linear backward model that attempted to reconstruct the fundamental waveform of an unrelated speech segment from the same female speaker. The noise models were computed using the same methodology we employed for determining the actual brainstem response, including the same cross-validation procedure and the same determination of the optimal regularization parameter per subject.

We then assessed whether the correct linear backward model outperformed the noise model, or the opposite, by comparing the correlation coefficients obtained on the ten-second segments through a two-tailed Wilcoxon signed rank test. The results of the statistical tests are indicated for each subject in Figure 1-A through asterisks: no asterisk is given when results are not significant (*p* > 0.05), one asterisk when results are significant (*,0.01 < *p* ≤ 0.05), two asterisks when significance is high (**, 0.001 < *p* ≤ 0.01), and three asterisks when significance is very high (***, *p* ≤ 0.001).

### Estimation of the neural response (forward model)

To gain further information about the neural origin of the response we also computed a linear forward model that estimated the EEG responses from the fundamental waveform. The coefficients of the forward model, as opposed to those of a backward model, allow for a neurobiological interpretation of their spatio-temporal characteristics (Haufe et al., 2014). The forward model relates the EEG recording *x_j_*(*t_n_*) at time *t_n_* to the fundamental waveform *y*(*t_n_* – *τ_k_*) as well as its Hilbert transform *y^h^*(*t_n_* – *τ_k_*) at a delay *τ_k_*:

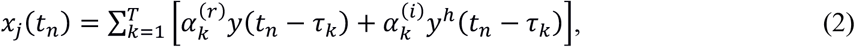

in which 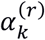 and 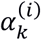 are the model’s real coefficients. They can be interpreted as real and imaginary parts of the complex coefficients 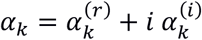. To investigate the temporal dynamics of the neural response, we considered a broader range of time lags than for the backward model. Specifically, we employed a set of *T*=201 possible delays *τ_k_* that ranged from −50 ms up to 150 ms with an increment of 1 ms. Although we did not expect a neural response at negative delays or at delays larger than 20 ms, we included those nevertheless to verify the absence of a significant response there. The model coefficients were estimated by concatenating the data from all subjects that showed a significant brainstem response to the speech signal as assessed earlier (generic or subject-averaged model) and using a regularised ridge regression as previously described.

As for the backward model, we made the Python code for computing the complex coefficients of the forward model available on Github as well (Kegler et al.).

### Significance of the auditory brainstem response

We sought to investigate at which latencies significant neural responses emerged. We therefore compared the obtained forward model to noise models. One thousand forward noise models were computed analogously to the forward model, except that the fundamental waveform of the actual speech signal was replaced with a fundamental waveform of an unrelated speech stimulus, from the same female speaker. We constructed these unrelated speech stimuli by randomly picking four parts, each with a duration of 2.5-minutes, from the eight parts that constituted the female speech material used in the competing speaker condition. This procedure was repeated to create 1,000 surrogate waveforms (out of all 1,680 possible combinations). We then employed a mass-univariate analysis to identify the significant time delays (Groppe et al., 2011). In particular, we computed the average magnitude of the responses over the EEG channels, yielding a single real time-varying function for the actual neural response and of the noise responses. We then pooled the values from the 1000 noise responses over the time lags to establish a single empirical null-distribution. We used this distribution to determine a critical value corresponding to a *p*-value of 0.05 to which the actual neural response was compared at each time lag from −50 ms to 150 ms (Bonferroni correction for multiple comparison).

In addition, we analysed the topography of the forward model at the peak latency *τ*_0_ of the average magnitude of the responses over the EEG channels. To this end, the forward noise models were used to build an empirical null distribution for each channel. For each noise model, the peak latency of the average magnitude was determined, and the magnitude of each channel’s response at this latency was used to establish the null distribution of that channel. Finally, the forward model at time *τ*_0_ was compared to the corresponding null empirical distribution at the respective channel at a significance level of *p* = 0.05, with FDR correction for multiple comparison over channels.

### Stimulus artifacts

We also computed the cross-correlation between the EEG responses to speech in quiet and the corresponding broad-band speech signal, with the purpose of checking for stimulus artifacts. To this end the speech stimulus was resampled from 44,100 Hz to 1,000 Hz, the sampling frequency of the EEG data. The cross-correlation functions were then analysed for statistically significant peaks at delays between −200 ms to 200 ms following the same procedure as described above for the forward model. Briefly, the cross-correlations were first averaged over subjects, and the absolute value of the resulting functions were then averaged over electrodes, yielding the average neural response as a function of latency. To establish a chance level, the same calculations were reproduced when replacing the speech stimulus by a different one from the same speaker. This procedure was repeated 1,000 times, yielding 1,000 noise responses. These stimuli were constructed as described above. These noise responses were pooled over time lags to build a single null distribution that was then used to assess the significance of the actual averaged neural responses as described above for the forward model (*p* = 0.05, Bonferroni-corrected for multiple comparison over time lags between −200 ms to 200 ms).

### Attentional modulation of the auditory brainstem response

To analyse the attentional modulation of the brainstem response to one of two competing speakers, we computed two pairs of backward models for each subject. The first pair of models reconstructed the fundamental frequency of the male voice while it was either attended (MA model) or ignored (MI model). The second pair of models reconstructed the fundamental waveform of the female voice when the subject attended it (FA model) or when the subject ignored it (FI model). The computation of the backward models, and the assessment of their performance, was done through five-fold cross-validation as explained above.

For each speaker, the performances of the attended and ignored models were then compared using a two-tailed Whitney-Mann rank test at the subject level. The results are indicated in Figure 4 through asterisks as described above. We further employed a two-tailed Wilcoxon signed-rank test to investigate whether the population-average ratios of the performances were, for each speaker, significantly different from unity. Finally, we used a two-tailed Wilcoxon signed-rank test to check if the population-average ratios obtained from the responses to the male voice and to the female voice were significantly different.

### Differences between brainstem responses to attended and to ignored speech

We sought to determine whether the difference in the brainstem response to attended and to ignored speech reflected merely a difference in the strength of the response, or if there were other as well. To this end, we compared the magnitudes and the phases of the complex coefficients of the forward model for an attended voice to those for an ignored voice. Because the forward models for the male and for the female voice reflected the different fundamental frequencies of both speakers, we performed this analysis separately for the male and for the female voice. Regarding the magnitude, we computed the ratio of the amplitude of the attended and of the unattended model, at the peak delay of their average amplitude (9 ms, for both the male and female voices). We then employed a two-tailed Wilcoxon sign-rank test to determine for which electrodes the ratio was significantly different from unity (*p* < 0.05, FDR-corrected for multiple comparison over electrodes). To compare the phase, we computed the phase difference between the attended and the ignored model at each electrode at this same peak latency. We considered the wrapped phase differences that were mapped to the range of [−π, π]. We then determined the statistical significance of the phase difference through the Rayleigh test for non-uniformity of circular data (*p* < 0.05, FDR-corrected for multiple comparison over electrodes). The Rayleigh test assesses the null hypothesis that the phase differences are uniformly distributed around the circle. However, it does not inform on the value of the phase differences. Therefore, we derived 95% confidence intervals for the mean phase difference by pooling the data across all electrodes that exhibited significant phase clustering. All circular statistics were performed using the Circular Statistics Toolbox for Matlab (Berens, 2009). Finally, we compared the latency of peak amplitude between the attended and ignored models using a Wilcoxon signed rank test.

In order to enable a direct comparison with our previous related work, we also computed the difference between the TRF at electrode CPz and the average TRF of the two mastoids to produce one dipolar response (Forte et al., 2017). CPz was selected due to its central location, similar to the one used in our previous study, and because it emerged in our present study as one of the central electrodes that displayed a significant response to speech in quiet (Figure 1-C). We then computed the ratio of this dipolar response between the attended and the ignored condition.

### Decoding of auditory attention

We investigated how attention could be decoded from short segments of neural data that were obtained in response to competing speakers. We first trained and assessed the performances of the two pairs of speaker-specific linear backward models (MA, MI, FA, FI, as described above) using five-fold cross-validation. For all the attention decoding procedures presented hereafter, the normalised regularisation coefficient of the backward models was fixed to the value that yielded the best reconstruction for speech in quiet, *λ_n_* = 10^−0.5^.

The testing fold was divided into testing segments with a duration of 0.5, 1, 2, 4, 8, 16 and 32 s. For each testing segment we therefore obtained four different correlation coefficients: the correlation coefficient r_MA_ between the fundamental waveform of the male speaker and its reconstruction based on the MA model, the correlation coefficient r_MI_ between the fundamental waveform of the male speaker and its reconstruction based on the MI model, as well as the correlation coefficients r_FA_ and r_FI_ between the fundamental waveform of the female speaker and its reconstruction based on the FA and FI model, respectively. The computed correlation coefficients were then employed to decode attention on each segment. We thereby explored two different avenues (Figure 6-A).

First, we based the decoding on the attended models MA and FA only. To this end, we compared the correlation coefficients from both models. If r_MA_ exceeded r_FA_ we concluded that the male speaker was attended, and otherwise that the female speaker was the focus of attention. Second, we considered the ignored models MI and FI only. If r_MI_ was larger than r_FI_ attention was decoded as having been directed at the female speaker, and *vice versa* if r_MI_ was smaller than r_FI_.

The decoding of attention using these two different methods was performed using all 64 EEG channels as well as based on three EEG channels only (vertex and mastoids: Cz, TP9, TP10). The decoding of attention based on the attended models was also performed using the fundamental waveform obtained by band-pass filtering.

We sought to compare the performance of the obtained attentional decoding to that of a random classifier. A random binary classifier can achieve a high accuracy by chance. This is especially true when the number of testing data is small, which in our case occurs when the duration of the testing segments is long. To account for this effect, we determined the 95% chance level, that is, the highest accuracy that a random classifier would achieve in at least 95% of cases. This 95% chance level was computed using a binomial distribution (Combrisson and Jerbi, 2015).

### Subject-independent attention decoding

In real-life situations, the decoding of auditory attention may be required for a subject for whom training data is not available. This situation requires to train a decoder on other people for whom training data is at hand, and to then apply it to the subject under consideration. We refer to such decoders as out-of-the-box models since, once trained on the data from a set of volunteers, they can be readily applied to other subjects. To assess how well these out-of-the-box models decode auditory attention, we trained linear backward models on the pooled data from all subjects and quantified their performances using a leave-one-subject-out cross-validation coupled with a five-fold cross-validation regarding the auditory stimuli (*i.e*. testing on data from a subject and from a part of the stimulus unused during training). To train the model, the testing data from all-but-one participants was concatenated and used to obtain the model coefficients. The unseen part of the data from the remaining subject was used to assess the performance of the model. In particular, we assessed the classifier that compared the performances of the MA and the FA model. Its classification accuracy was evaluated as described above.

### Speaker-averaged attention decoding

We also wondered how well selective attention could be decoded from the brainstem response if the specific models of the brainstem responses to the individual voices were not available. We therefore followed a similar analysis as used for decoding auditory attention based on the speech envelope (O’Sullivan et al., 2014). For each subject, we computed a single backward model for an attended voice, irrespective if it was the male or the female one. This model was accordingly trained on the data from both the condition when the subject attended the male voice and the condition when they listened to the female speaker. The male fundamental waveform was used as the reconstruction target when the male speaker was attended, and the female fundamental waveform was the target when the female voice was attended. An equal proportion of data from each attention condition was included in each cross-validation fold. To determine the focus of attention, we then considered short testing segments as described above. For each testing segment we computed the correlation coefficient between the reconstructed fundamental waveform and the actual ones of the two speakers. If the reconstruction matched the fundamental waveform of the male speaker more closely than that of the female one, we concluded that the subject had attended the male speaker. Otherwise we determined that the focus of attention was on the female voice. The performance of the classifier was evaluated as described above.

## Results

### Response to a single speaker

We first measured neural responses to a single non-repetitive speech signal from 64-channel EEG. We employed empirical mode decomposition to obtain a fundamental waveform from the speech signal (Forte et al., 2017), and linear regression with regularization to reconstruct the fundamental waveform from the multi-channel EEG data for each individual subject (linear backward model, Methods). The performance of the reconstruction was assessed through the mean Pearson’s correlation coefficients over ten-second segments of the reconstructed fundamental waveform to the actual one (Figure 1-A).

We verified that the linear backward models did extract a significant brainstem response to speech. To this end we also constructed models based on the fundamental waveform of unrelated speech signals from the neural data. For almost all subjects that we assessed (15 out of 18), the model that reconstructed the actual fundamental waveform significantly outperformed the one that attempted to reconstruct an unrelated fundamental waveform, showing that the former was able to extract a meaningful brainstem response (Figure 1-A, two-tailed Wilcoxon signed-rank test).

To investigate the spatio-temporal characteristics of the brainstem response we computed a generic linear forward model that estimated the EEG recordings from the fundamental waveform using the data from all the subjects that yielded significant reconstructions in the previous test presented in Figure 1-A (Methods). The average over channels of the magnitude of the obtained complex coefficients peaked at 8 ms, and only the latencies around this peak (3 to 14 ms) yielded statistically-significant neural responses (Figure 1-B). This finding demonstrated the subcortical origin of the neural activity and was in agreement with previous recordings of speech-evoked brainstem responses (Skoe and Kraus, 2010; Reichenbach et al., 2016; Forte et al., 2017; Maddox and Lee, 2018). The magnitude of the coefficients at that latency showed major contributions from the mastoids as well as moderate contributions from the central scalp areas (Figure 1-C). Both the mastoid channels as well as the channels near the midline of the scalp yielded significant responses. The coefficients at the central area were approximately in antiphase to those near the mastoids, reflecting the direction of the brainstem’s dipole sources (Figure 1-D).

We also computed linear forward models for single subjects (Figure 2). We find that they yielded peak responses at similar latencies, and showed similar topographies, although these were noisier than the ones obtained from the average over all subjects.

### Absence of stimulation artifacts

To determine if stimulus artifacts were present in the recordings, we computed a cross-correlation between the EEG data and the broadband speech signal. Broadband speech elicits neural responses from the brainstem to the cortex, at latencies ranging from 5 ms to a few hundred ms (Maddox and Lee, 2018). A stimulus artifact would arise, in contrast, instantaneously, at a delay of −1 ms. This delay reflects the fact that, in our analysis, we compensated for the earphone’s 1 ms delay of delivering the sound to the ears. The responses that we recorded contained, however, only significant contributions between 9 and 12 ms delays, firmly in the range of subcortical neural activity (Figure 3). We could accordingly not detect stimulus artifacts in our EEG recordings.

### Attentional modulation of the response to competing speakers

We then investigated how attention modulates the brainstem response. Following a classic auditory attention paradigm we presented subjects with a male and a female voice diotically and simultaneously, instructing them to attend to either the male or the female speaker, while recording their neural activity from 64-channel EEG (Ding and Simon, 2012; Forte et al., 2017). For each subject, we computed four linear backward models. The first model, MA, reconstructed the fundamental waveform of the male voice when the subject attended to it. The second model, MI, reconstructed the fundamental waveform of the male speaker when the subject ignored it. Analogously, a third and fourth model, FA and FI, reconstructed the fundamental waveform of the female voice when it was attended or ignored, respectively. We observed that the performance of the two models that reconstructed the fundamental waveform of a speaker when they were attended was, in most subjects, significantly better than that of the corresponding model for the ignored voice (Figure 4, two-tailed Whitney-Mann rank test). The average ratio between the reconstruction performance of the model for the attended male voice to that for the ignored male voice was 1.22, significantly larger than one (Z(17) = 7, *p* < 0.001, two-tailed Wilcoxon signed-rank test). The ratio was 1.15 in the case of the female voice, which was significantly above one as well (Z(17) = 38, *p* = 0.039, two-tailed Wilcoxon signed-rank test). The two ratios did not differ significantly (Z(17) = 69, *p* = 0.47, two-tailed Wilcoxon signed-rank test). The better reconstruction performance of the fundamental waveform of an attended speech signal demonstrates the attentional modulation of the brainstem response to speech that we described previously (Forte et al., 2017).

We wondered if the difference between the attended and the ignored brainstem response reflected merely a difference in the strength of the response, or if there were other differences as well. To investigate the nature of these differences, we compared the coefficients of the attended forward models to those of the ignored models, at the peak delay of their average amplitude (9 ms). We found that the ratio of the magnitude of the coefficients did not differ statistically from unity, neither for the male nor for the female voice (Figure 5-A,C; Wilcoxon sign-rank test, FDR correction for multiple comparison over electrodes). However, we found a statistically significant clustering of phase differences between the attended and the ignored models at several electrodes near the midline as well as near the mastoids (Figure 5-B,D; Rayleigh test for non-uniformity of circular data, FDR correction for multiple comparison over electrodes). For the male voice, the mean phase difference was found to be −0.51 π (95 % confidence interval: [−0.56 π; −0.47 π]), while it was −0.12 π for the female voice (95 % confidence interval: [−0.17 π; −0.08 π]). This shows that the ignored models were not merely a scaled version of the attended models, but that the brainstem response to ignored speech occurred at a different phase from that to attended speech.

Due to the range of frequencies that constitute the fundamental waveform, the phase shift between the attended and the ignored models did not equate to a consistent temporal shift. We did indeed not find a statistically-significant difference in the timing between the peak amplitude of the attended and the ignored models across the different subjects, for the male or female voice (*p* = 0.17 and *p* = 0.69 respectively, two-tailed Wilcoxon signed rank test).

To facilitate comparison with previous work we also computed the difference of the mastoid electrodes and the electrode at CPz, yielding a dipolar response (Forte et al., 2017). We found that the response’s ratio between the attended and ignored condition was significantly greater than unity, for both the male and female voices (p = 0.016, and p = 0.003 respectively, Wilcoxon sign-rank test).

### Decoding of auditory attention

Having verified the attentional modulation of the brainstem response to speech using high-density EEG recordings and linear backward models, we sought to investigate whether this approach could be used to decode auditory attention. We expected the focus of attention to emerge, for instance, from the difference in the performances of the models MA and FA. This difference should typically be positive when the subject attended to the male voice and be negative otherwise (Figure 6-A). Similarly, attention could potentially be decoded from the difference of the reconstruction performance of the models FI and MI. A subject’s attention to the male voice should mostly lead to a positive difference, and a focus on the female voice to a negative difference.

We tested the accuracy of the decoding on samples of a duration that varied from half a second to over 30 seconds (Figure 6-B). The averaged decoding accuracy based on the attended models (MA, FA) remained significantly above chance even for very short samples that lasted only half a second. It was, for instance, 59% and 69% for two-second and sixteen-second samples, respectively. In contrast, the models MI and FI by themselves did not allow for a decoding of the attentional focus with an accuracy that was better than chance. In the following we therefore discuss decoding obtained from the attended models only.

Practical applications of the decoding of auditory attention benefit from a small number of required recording channels. We therefore investigated how well the developed decoding works if the linear backward models use only three EEG channels, the left and right mastoid as well as the central channel Cz. Strikingly, the subject-averaged decoding accuracy was barely smaller than that of the 64-channel model; for instance, it remained at 69% for a sixteen-second sample when the classifier based on the attended models was used (Figure 6-C).

Both for the 64-channel as well as for the 3-channel decoding we observed variation in the decoding accuracy from subject to subject (Figure 7-A). For a duration of 16 s, for instance, some subjects showed decoding accuracy close to 90%, whereas other subjects exhibited significantly lower decoding accuracies that did not exceed the change level. However, even for short testing segments and for the majority of subjects, the decoding remained above chance level. We note in addition that the subjects that did not allow for significant decoding include those for whom we did not obtain significant brainstem responses to speech in quiet (Figure1-A).

Because of the complexity of empirical mode decomposition (EMD), the computation of the fundamental waveform through this method cannot typically be performed online. We therefore wondered if attention could be decoded based on a similar waveform obtained through band-pass filtering the audio signal in the range of the fundamental frequency. Band-pass filtering is indeed a comparatively simple operation that can run in real time. We found that decoding based on the band-pass filtered audio has a similar accuracy as the one based on the waveform obtained from EMD, which is encouraging for real-time applications (Figure 7-B).

Real-world settings will often feature voices that have not been encountered before and for which no speaker-specific model of the brainstem response is available. In an attempt to generalise our results, we computed a speaker-averaged backward model for *any* attended speaker, irrespective of whether it was the male or the female one. We then decoded attention from the performance of this speaker-averaged model in reconstructing the fundamental waveform of either the male of the female speaker. The averaged decoding accuracies that we obtained were slightly lower than those from the speaker-specific models but were above chance level for durations down to 0.5 s (Figure 7-C).

The decoding described above utilized linear backward models that were subject specific and hence required prior training from EEG recordings for each individual. Such subject-specific training may, however, not always be available. We thus assessed the performance of a linear backward model that was trained on the whole population of subjects, and thus represented an average model that could be used out-of-the-box to decode attention. As expected, the decoding accuracies were then lower than those for the subject-specific models. While the decoders based on the attended models with all 64-channels remained above the chance level for all the tested durations, the 3-channel setup yielded worse performance only slightly exceeding the chance level for all but the longest duration. For duration of 16 s, for instance, the 64-channel setup yielded 65% accuracy, while the 3-channel only 63% (Figure 7-D). Although the accuracy of this decoding when averaged across subjects was not very high, we note that this average was significantly reduced by a few subjects that showed particularly poor accuracies of around 50%, reflecting poor brainstem recordings from these subjects. The majority of the subjects, in contrast, yielded decoding accuracies that exceeded the chance level.

## Discussion

We showed that the brainstem response to the fundamental frequency of speech can be measured reliably from high-density EEG recordings in most subjects through a statistical modelling approach. The response is most evident in the differences between the electrodes near the mastoids and those close to the vertex, in agreement with the dipolar structure of scalp-recorded auditory brainstem activity (Ono et al., 1984; Grandori, 1986; Norrix and Glattke, 1996; Bidelman, 2015). Moreover, the response latency of 8 ms evidenced a subcortical origin.

The frequency-following response (FFR) to simpler acoustic signals such as long vowels has recently been found in an MEG study to contain cortical contributions (Coffey et al., 2016). However, when measured through EEG, the cortical contributions emerge earliest at a latency of 20 ms, are smaller than the subcortical ones, and mostly apparent for frequencies up to about 100 Hz (Bidelman, 2018). The response to the fundamental frequency of running speech that we have measured here does not show a measurable signal at latencies longer than 14 ms and was recorded in response to a fundamental waveform high-pass filtered above 100 Hz. While contributions from cortical structures cannot be entirely ruled out, we did not observe any within our measurement accuracy.

When subjects switched attention from one to another of two competing speakers, we found that the fundamental frequency of each speaker was better encoded in the brainstem response when that speaker was attended rather than ignored. These results align with those that we obtained previously from different recording equipment and with a different analysis procedure that did not involve statistical modelling and that did not address attention decoding (Forte et al., 2017). Here we found, however, that the ratio of the attended to the ignored temporal response functions, as obtained from the forward models, did not differ significantly between the male and the female voice. Indeed, although the scalp maps that we derived largely showed a larger response to the attended than to the ignored speaker (Figure 5-A, C), the modulation was not statistically significant. This presumably reflected the inclusion of all electrodes in the forward model, including many electrodes that displayed a poor signal-to-noise ratio. The backward models, in contrast, employed a weighting of the contribution from each electrode which boosted those with a large signal-to-noise ratio and thus led to a more significant result. To further investigate this issue, we also computed the response at a single channel that was obtained as the difference between the electrodes at the mastoids and at CPz, mimicking our previous bipolar recordings (Forte et al., 2017). The amplitude of this response was significantly modulated by selective attention, in agreement with our previous results.

The modelling work that we developed here allowed us to further investigate the origin of the difference in the brainstem response to attended and to ignored speech. We thereby found a significant difference between the phases of the response to attended versus ignored speech. Such a phase shift could in principle emerge from a difference in latency between the attended and ignored model. However, we found no statistically significant difference in peak latency of the attended and ignored responses. The phase shift might instead signify different relative contributions of different parts of the brainstem to the scalp-recorded response. The different values of the phase shift that we obtained for the male and female voice may reflect the differences in the fundamental frequencies of both stimuli.

Most importantly, we developed a procedure to decode the attentional focus of a subject to speech based on her or his brainstem response as measured from as few as three recording channels. This will enable the future characterization and investigation of the subcortical mechanisms through which the brain solves the cocktail party problem. Potential practical applications include brain computer interfaces, such as neuro-steered auditory prostheses, as well as clinical assessments of suprathreshold hearing impairments that cannot be identified from pure-tone audiometry. Any of these applications will benefit from a decoding method that is fast and requires only a small number of recording channels.

We showed that the best decoding is achieved when linear models that relate the neural recording to the speech signal are computed for each subject individually. Such subject-specific models may cause difficulty in practice as sufficient training data per subject may not always be obtainable. The out-of-the-box models reflect the generalized version of the models obtained from the data pooled over many subjects and can be readily applied to other subjects for which no training data is available. We have shown that while the decoding performance of the out-of-the-box models is below those of the subject-specific models, the average decoding accuracy still exceeds the noise level for the high-density EEG setup. This suggests a consistency of the brainstem responses to speech across the participants. We also note that the out-of-the box models were fitted using the data from all subjects, including those that did not yield a significant reconstruction of the fundamental waveform in the speech-in-quiet condition.

Potential real-world applications will also often require the decoding of attention to a speaker that has not been encountered before. As an important step in this direction, we showed that speaker-averaged models that are trained on both attended speech signals, thereby computing an attended model that was averaged over the different voices, still performed well and allowed to decode attention. Future work could investigate how well these models generalise to speakers for which no training data is available.

Another important feature for real-time attention decoding is that the whole computational pipeline – from the processing of the audio signal to the computation of reconstructed waveforms and the attention decoding – can run online. Our reconstruction of the fundamental waveform through a backward model, the assessment of its performance as well as the subsequent attention decoding were all based on linear operations that can easily run in real time. However, the EMD that we employed for the computation of the fundamental waveform comes with large computational costs. We therefore explored how a computationally much simpler operation, band-pass filtering of the audio signal, performed regarding the decoding of attention. Promisingly we found that this method still allowed to decode attention from very short segments of data, evidencing the potential for real-time decoding. While two band-pass filters with different corner frequencies were applied to the male and female voice, this approach could be extended to use filterbanks or use online pitch estimation algorithms.

The decoding procedure that we developed relies on the correlation between the reconstructed fundamental waveform from the brainstem response and the actual fundamental waveform of the speech signal. The obtained correlation coefficients are small, typically between 0.05 and 0.1 (Figure 1-A, Figure 2). Cortical responses allow to reconstruct the brainstem response from EEG recordings and yield somewhat higher correlation coefficients. However, the attentional decoding based on the brainstem responses that we show here is comparable to the decoding based on the reconstructed speech envelope, obtained from 64 EEG channels. A 16-s trial, for instance, yields an average decoding accuracy of about 69% when based on the fundamental waveform, which is similar to the corresponding decoding accuracy that was reported in several previous studies (O’Sullivan et al., 2014; Biesmans et al., 2016; Bleichner et al., 2016). We attribute this similarity of the attention decoding accuracies to the rapidness of the brainstem response: because the brainstem response to speech occurs at the fundamental frequency of a voice, it is ten- to hundredfold faster than the cortical response to the speech envelope. This rapidness appears to compensate for the smaller magnitude of the response.

Although brainstem responses and cortical responses allow for similarly efficient attention decoding when high-density EEG is available, the decoding based on the brainstem response to speech may have advantages when only a few channels are available. The accuracy of attention decoding based on the speech envelope drops indeed below 80% for a trial of at least 20 seconds when relying on subject-specific five-electrode montages (Mirkovic et al., 2015; Fuglsang et al., 2017). Similarly, the attention decoding based on the brainstem response that we have developed here achieves an averaged accuracy of 69% when based on three electrodes (TP9, TP10 and Cz) and on 16 seconds of data, and reaches 72% when 32 seconds of data are available (Figure 5-B). This good decoding performance from a few EEG channels may be due to the effective capturing of the brainstem response by sparse montages, as well as due to a consistent dipole orientation across subjects (Dale and Sereno, 1993). Importantly, we employed only band-pass filtering as a pre-processing step for the EEG data. The simplicity of this attention decoding method and its good accuracy when based on a few EEG channels may make this method attractive for practical applications.

The mixed-speaker stimuli that we employed were obtained by superimposing two speech signals, and our decoding was based on the knowledge of these separate voices. The individual components of a complex acoustic scene are, however, in general not available and need to be estimated from the acoustic mixture. The application of our method for decoding attention to steer an auditory prosthesis towards an attended voice, for instance, will thus require to first segregate the different voices that are present in the acoustic space, and to then determine the focus of the user’s attention. The segregation of the different individual speakers may be achieved through multi-microphone arrays together with methods such as beamforming (Gannot et al., 2001) or non-negative blind source separation (Van Eyndhoven et al., 2017).

Certain applications may, however, not require the separation of the individual voices from an acoustic mixture but have them already available. Many locked-in patients, for instance, cannot communicate overtly, not even through eye motion (Giacino et al., 2002). Current brain-computer interfaces for them are mostly based on the P300 response, an evoked cortical potential that arises 300 ms after the occurrence of an oddball stimulus. It is typically elicited through visual or through sound stimuli and requires a few seconds to achieve a single binary response (Piccione et al., 2006; Nijboer et al., 2008; Schreuder et al., 2011). A brain-computer interface based on auditory attention, in contrast, could present a mixture of two auditory streams to the patient. The patient could then answer a question with yes or no through attending to a particular stream. Because the stimuli are merely used as a locus of attention, they would be available individually beforehand, and could be engineered to enhance decoding speed. Similarly, clinical assessments of the brainstem response to speech and its modulation through selective attention can employ predefined acoustic mixtures.

The decoding that we have described here is based on linear backward models that reconstruct the fundamental waveform of the speech signal from the EEG recordings. This method determined the brainstem response to the voiced parts of speech, and in particular to its pitch, but did not measure the brainstem response to the voiceless speech components (Maddox and Lee, 2018). Improved performance may be obtained through canonical correlation analysis that relates the neural recording to more speech features in an optimized space (de Cheveigné et al., 2018) or through an artificial neural network that is able to extract highly nonlinear relations between the two datasets (Yang et al., 2015).

Finally, decoding of auditory attention could leverage both cortical and sub-cortical responses as they can be obtained from the same EEG recordings. The framework for attentional decoding based on the brainstem response to running speech presented here could be readily extended to include cortical responses to the speech envelope, which could boost the overall decoding accuracy. Moreover, measuring both subcortical and cortical responses to speech from the same EEG data will be useful for fundamental auditory research and clinical assessment of hearing impairments.

## Acknowledgement

This research was supported by EPSRC grants EP/M026728/1 and EP/R032602/1 to T.R., by the Royal British Legion Centre for Blast Injury Studies, by Wellcome Trust grant 108295/Z/15/Z, as well as in part by the National Science Foundation under Grant No. NSF PHY-1125915. We are grateful to the Imperial College High Performance Computing Service (doi: 10.14469/hpc/2232). We thank Steve Bell, Karolina Kluk-de Kort, Patrick Naylor, David Simpson, Alain de Cheveigné and Malcolm Slaney for discussion.

## Figure caption

**Figure 1**. The brainstem response to natural speech detected from high-density EEG recordings using complex linear models. (**A**) The performance of the linear backward model is quantified through the Pearson’s correlation coefficient of the reconstructed fundamental waveform and the actual one. For each subject the presented result is the averaged correlation coefficient obtained from 10-seconds long segments of the EEG and the fundamental waveform (white bars). In almost all subjects, the performance is significantly better than that of a model estimating the noise-level reconstructions. Subjects have been ordered by increasing performances. (**B**) The channel-averaged magnitude of the complex coefficients of the generic forward model obtained from the pooled data from all the participants that yielded significant reconstructions, peaks at a latency of 8 ms. Only latencies ranging from 3 to 14 ms yield a statistically-significant response (black bar, *p* < 0.05, Bonferroni correction), as compared to noise models. (**C**) At the delay of 8 ms, a significant neural response emerges from the mastoid channels as well as from the channels near the midline (white disks, *p* < 0.05, FDR correction, population average). (**D**) The phase of the complex coefficients at the delay of 8 ms shows a phase difference of around *π* between the temporal areas and the central one (population average).

**Figure 2**. Brainstem responses to speech from two single subjects. The top row shows the brainstem response from subject 9 that yielded the median reconstruction of its brainstem response to speech (Figure 1). The bottom row presents the results from subject 18 that had the best reconstruction of the brainstem response to speech. (**A**) The channel-averaged magnitude of the complex coefficients of the forward model peaks at a latency of 9 ms (subject 9) and 10 ms (subject 18). (**B**) The topographic maps of the coefficient magnitudes at the peak latency are consistent with those of the generic model, although more noisy in the case of subject 9. Channels located at the mastoids show the highest magnitudes. (**C**) The phase of the complex coefficients at the peak latency. The phases differ between the two subjects since they have been taken at different latencies (9 and 10 ms, respectively). Consistent with the generic model, the topographic plots show a phase difference of around *π* between the temporal areas and the central area.

**Figure 3**. Absence of stimulus artifacts. Magnitude of the cross-correlation between the EEG data and the broadband speech stimulus averaged over channels and participants. The only time lags for which the cross correlation is significantly greater than the estimated noise floor are between 9 - 12 ms. In particular, the model shows no significant response at the delay of −1 ms, the delay of the earphones, evidencing the absence of stimulus artifacts.

**Figure 4**. Attentional modulation of the auditory brainstem response to natural speech. The order of the subjects is as in Figure 1A. (**A**) The performance of the linear backward model for the male voice is better when the male speaker is attended (black) then when he is ignored (red). The two performances differ significantly in most subjects, and so do the two average performances (avg). The average ratio between the two performances is 1.22 and is significantly larger than one (*p* = 0.01). (**B**) The performance of the linear backward model that reconstructs the fundamental waveform of the attended female voice is likewise significantly better than that for the ignored female voice in most subjects, as well as on average (avg). The average ratio of the two performances is 1.15 and is significantly larger than one (*p* = 0.039). The ratios for the male and female voices do not differ significantly (*p* = 0.47).

**Figure 5**. Differences in the brainstem response to attended and to ignored speech. (**A, C**) The subject-averaged ratio of the magnitude of the complex coefficients of the attended forward model to those of the ignored model, at the average peak latency of 9 ms. None of these ratios are statistically different from unity (FDR correction). (**B, D**) The subject-averaged phase difference between the coefficients of the attended and the ignored forward models, at the average peak latency of 9 ms. Channels close to the midline as well as at channels near the mastoids yielded a significant phase difference (*p* < 0.05, FDR correction). The male models exhibit a phase difference of −0.51 π (95 % CI: [−0.56 π; −0.47 π]), while the female model phase difference is −0.12 π (95 % CI: [−0.17 π; −0.08 π]).

**Figure 6**. Decoding of auditory attention. (**A**) Testing data of a duration of 32 s that were obtained from a subject listening to the male speaker (black) can potentially be discriminated from those obtained when a subject listened to the female voice (red) through the performances *r* from four linear backward models (MA, MI, FA, FI; Methods). The classification can employ the difference in the performances between the models MA and FA (green) or the difference between the models FI and MI (orange). (**B**) The subject-averaged decoding accuracy obtained from the models MA and FA reaches 73% at a duration of 32 seconds and remains above chance level (grey) for very short durations of 500 ms. Decoding based on the models FI and MI remains below chance level (average over all subjects). (**C**) Employing only three recording channels to decode attention reduces the performance of the classifiers only slightly, if at all.

**Figure 7**. Different types of attention decoding and intra-subject variability. The two rows of panels correspond to the 64-channel and to the 3-channel decoders, respectively. (**A**) The attention decoding accuracies from the speaker-specific models achieved per individual subject (coloured lines, consistent across panels) varies by up to approximately 50% around the average (bold black line). However, for each individual subject the decoding based on 64 channels (top) is similar to that achieved from three channels (bottom). Here, the decoding is based on the difference between the attended models (same data as presented on the population level in Figure 6-B,C by the green lines). (**B**) Instead of using empirical mode decomposition (EMD), a fundamental waveform can be estimated by band-pass filtering the speech signal, which can be implemented in an online fashion. Attention decoding based on the band-pass filtered audio achieves a similar performance as the one based on the EMD. (**C**) Attention can be efficiently decoded using a single attended model for both speakers as well. (**D**) The use of the out-of-the-box backward models for reconstructing the fundamental waveforms, leads to reduced, yet better than chance, decoding accuracies for most subjects.

